# Resistance and Tolerance to Root Herbivory in Maize were Mediated by Domestication, Spread, and Breeding

**DOI:** 10.1101/751982

**Authors:** Ana A. Fontes-Puebla, Julio S. Bernal

**Author notes:** **Correspondence:** Julio S. Bernal.

## Abstract

Plants may defend against herbivory and disease through various means. Plant defensive strategies against herbivores include resistance and tolerance, which may have metabolic costs that affect plant growth and reproduction. Thus, expression of these strategies may be mediated by a variety of factors, such as resource availability, herbivory pressure, and plant genetic variation, among others. Additionally, artificial selection by farmers and systematic breeding by scientists may mediate the expression of resistance and tolerance in crop plants. In this study, we tested whether maize defense against Western corn rootworm (WCR) was mediated by the crop’s domestication, spread, and modern breeding. We expected to find a trend of decreasing resistance to WCR with maize domestication, spread, and breeding, and a trend of increasing tolerance with decreasing resistance. To test our expectations, we compared resistance and tolerance among four *Zea* plants spanning those processes: Balsas teosinte, Mexican landrace maize, US landrace maize, and US inbred maize. We measured performance of WCR larvae as a proxy for plant resistance, and plant growth as affected by WCR feeding as a proxy for plant tolerance. Our results showed that domestication and spread decreased maize resistance to WCR, as expected, whereas breeding increased maize resistance to WCR, contrary to expected. Our results also showed that maize resistance and tolerance to WCR are negatively correlated, as expected. We discussed our findings in relation to ecological-evolutionary hypotheses seeking to explain defense strategy evolution in the contexts of plant resistance-productivity trade-offs, plant tolerance-resistance trade-offs, and varying resource availability vis-à-vis plant physiological stress and herbivory pressure. Finally, we suggested that defense strategy evolution in maize, from domestication to the present, is predicted by those ecological-evolutionary hypotheses.

## INTRODUCTION

Though sessile, plants are not defenseless organisms incapable of escaping their enemies, and generally employ various means for defending themselves against herbivory and disease. When directed against herbivory, such defensive means include physical and chemical defenses, the ability to manipulate primary metabolite allocation to reduce herbivore fitness, and tolerance, which are important mediators of plant reproductive success (Zhou et al., 2015; Zust and Agrawal, 2017). Broadly, plant defensive strategies include resistance and tolerance. Resistance relies on direct (physical and chemical) and indirect (e.g., natural enemies, phenology) defenses, while tolerance involves compensatory growth, increased photosynthesis, and other responses that allow plants to reproduce without selecting for herbivore resistance and at no net metabolic cost (Painter, 1951; Strauss and Agrawal, 1999; Boege and Marquis, 2005; Schoonhoven et al., 2005; Stout, 2013). Generally, plant investment in defense seems to depend on resource availability, herbivore pressure, and genetic diversity (Hahn and Maron, 2016; Zust and Agrawal, 2017).

Whether below- or aboveground, defense against herbivores may be costly to both wild and cultivated plants. Generally, limited metabolic resources are distributed among multiple, competing processes, including defense (e.g., resistance) and productivity, (i.e. growth and reproduction). Defenses against herbivores may be constitutive, which are continuously present, or induced, which are summoned in response to herbivory. Subjected to herbivory, plants may allocate resources to defense responses accordingly, while other processes, such as reproduction (e.g., production of flowers, fruits, seeds), may be allocated fewer resources (Bazzaz et al., 1987; Rodriguez-Saona et al., 2011; Zust and Agrawal, 2017). However, in cultivated plants more resources tend to be allocated toward productivity than defense. For example, breeding for productivity and quality compromised defenses against herbivores in cranberries, so that herbivore performance was enhanced and constitutive and induced defenses were reduced on domesticated compared to wild cranberries (Rodriguez-Saona et al. (2011)). Also, in a study on *Zea* L. plants, maize wild relatives (*Zea mays* sspp.) were found to be better defended against herbivores, but had lower productivity, compared to modern cultivars of maize (*Zea mays mays* L.), which were poorly defended and had high productivity (Rosenthal and Dirzo, 1997). Interestingly, landrace maize, a form intermediate between maize wild relatives and modern maizes, showed intermediate defense and productivity. Overall, the study’s results supported a hypothesis positing that herbivore resistance in maize decreased with domestication and improvement for yield (Rosenthal and Dirzo, 1997).

Domestication, spread, and breeding are processes that can mediate crop evolution, including herbivore defense evolution. Accordingly, domestication modified interactions between crops and insects so that they differ substantially from those between crop wild ancestors and their herbivores (Macfadyen and Bohan, 2010; Chen et al., 2015a; Wang et al., 2018). For instance, following the initial domestication of maize ca. 9000 years before present (YBP) (Matsuoka et al., 2002), the sap-sucking herbivore *Dalbulus maidis* (Delong and Wolcott) became a pest as the crop’s defenses were weakened and as its distribution expanded from the Mexican subtropical lowlands to the temperate highlands and beyond (Medina et al., 2012; Bernal et al., 2017). As crops spread, they commonly face novel environmental variables, which may reshape plant-insect interactions (Baker, 1972; Erb et al., 2011; Meyer et al., 2012; Chen, 2016; Turcotte et al., 2017). Indeed, diverging climatic conditions, less competition, genetic drift associated with dispersal, among other variables, have been shown to produce changes in herbivory resistance in a variety of plants and crops (Rasmann et al., 2005; Zangerl and Berenbaum, 2005; Agrawal et al., 2012; Züst et al., 2012). Systematic breeding, along with geographical spread, also affects crop traits, including herbivore defenses. For example, maize underwent natural and artificial selection as it spread into new environments following its domestication (van Heerwaarden et al., 2012; Swarts et al., 2017; Kistler et al., 2018), and was subjected to systematic artificial selection (i.e. breeding) mainly for yield when agriculture was intensified in the 20^th^ century (Troyer, 1999; Whitehead et al., 2017). Such selection shaped maize’s herbivore defenses (Bellota et al., 2013; Davila-Flores et al., 2013; de Lange et al., 2014; Maag et al., 2015; Chinchilla-Ramírez et al., 2017). Moreover, enhanced plant growth in the face of novel herbivory pressure may lead to tolerance evolution, as posited under the resource availability hypothesis, which predicts that fast-growing plants in resource-rich environments, such as crop plants, may be selected to favor herbivory tolerance, at the expense of resistance (Rosenthal and Dirzo, 1997; Zou et al., 2007; Agrawal et al., 2010).

Crop plants can become hosts for herbivores as a consequence of domestication, spread to new environments, and breeding for high yield, as noted previously (Chen et al., 2015a; Chen, 2016; Chen and Schoville, 2018). After maize’s spread from the central Mexican highlands to North America, the oligophagous, root-chewing insect Western corn rootworm (WCR) (*Diabrotica virgifera virgifera* Le Conte) shifted to maize from an unknown ancestral Poeaceae host to later become a pest (Lombaert et al., 2017). WCR likely spread with maize from northern Mexico to southwestern United States (ca. 1500 CE) as maize became a significant crop and part of the Native American diet (Merrill et al., 2009; da Fonseca et al., 2015; Lombaert et al., 2017; Smith et al., 2017). WCR prefers maize over other hosts, which may be due to the crop plant’s comparatively weakened resistance against herbivory and greater nutritional value (de Lange et al., 2014; Bernal and Medina, 2018). Additionally, maize tolerance to WCR may have evolved as the crop faced less competition and non-native herbivory after its spread (Buckler and Stevens, 2006; Hahn and Maron, 2016; Robert et al., 2017). Currently, WCR is found in northern Mexico, USA, and Europe (Branson and Krysan, 1981; Gerdes et al., 1993; Gray et al., 2009). The economic damage that this herbivore can cause varies, e.g., economic losses attributed to WCR may exceed US$1B yearly in the USA (Gray et al., 2009), while in Europe they are estimated at €472 million per year (Wesseler and Fall, 2010).

Trade-offs between productivity (growth and reproduction) and herbivore resistance and between herbivore resistance and tolerance are at the base of hypotheses positing that with plant domestication and improvement for yield a crop’s resistance will suffer compared to that of its wild ancestor, and that tolerance increases as resistance decreases (Hahn and Maron, 2016). Indeed, prior studies comparing the defense responses of maize wild ancestors and maize exposed to different herbivores showed resistance de-escalations with domestication, spread, and breeding (Bellota et al., 2013; Szczepaniec et al., 2013; Bernal et al., 2015; Maag et al., 2015; Chinchilla-Ramírez et al., 2017), as well as increasing tolerance with spread (Zou et al., 2007). In this study, we tested whether maize defense against WCR was mediated by the crop’s domestication, spread, and breeding. To that end, we compared resistance and tolerance among four *Zea* plant types spanning those processes: Balsas teosinte (*Zea mays* L. spp. *parviglumis* Iltis and Doebley), Mexican maize landraces, USA maize landraces, and USA maize breeding lines. Each *Zea* plant type was represented by three accessions. The effects of domestication were assessed by comparing resistance and tolerance levels between Balsas teosintes and Mexican maize landraces; the effects of northward spread were assessed by comparing between Mexican landraces and US landraces, and; the effects of breeding were assessed by comparing between US landraces and US inbred lines. Specifically, we measured (i) performance of WCR larvae as a proxy for resistance, and (ii) plant growth as affected by WCR feeding as a proxy for tolerance. Overall, we expected to find decreasing resistance to WCR with maize domestication, spread and breeding, and increasing tolerance with decreasing resistance. We discussed our results in the context of plant resistance and tolerance evolution, as mediated by artificial and natural selection, geographical spread, and systematic breeding. Specifically, we discussed our findings in relation to ecological-evolutionary hypotheses seeking to explain defense strategy evolution in the contexts of plant resistance-productivity trade-offs, plant tolerance-resistance trade-offs, and varying resource availability vis-à-vis plant physiological stress and herbivory pressure.

## MATERIALS AND METHODS

### Plants and Insects

Four plant types belonging to the *Zea* genus were tested: Balsas teosinte, Mexican landraces, US landraces and US inbred lines (Table 1). These plant types were selected to represent the evolution of maize from its wild ancestor through the processes of domestication, spread, and breeding (Troyer, 1999; Matsuoka et al., 2002; Labate et al., 2003; Lombaert et al., 2017). Specifically, (i) Balsas teosinte is the immediate ancestor of maize, thus represented maize in its wild state, prior to domestication; (ii) Mexican landraces were included as descendants of Balsas teosinte, and served to assess the effects of domestication and the crop’s early upland spread; (iii) US landraces were included as descendants of Mexican landraces, and used to assess the effects of the crop’s spread to North America, and; (iv) US inbred lines were included as descendants of US landraces, and used to assess the effects of modern breeding. Three accessions were chosen as representatives of each of the plant types: “El Cuyotomate,” “Talpitita,” and “El Rodeo” for Balsas teosinte; Palomero Toluqueño, Chalqueño, and Cacahuacintle for Mexican landraces; Lancaster Sure Crop, Reid Yellow Dent, and Gourdseed for US landraces, and; MO17, B73, and W438 for US inbred lines (Table 1). The teosinte seeds were collected from subtropical lowland locations in Jalisco state, Mexico, whereas the Mexican landraces are grown in the central Mexican highlands. These landraces are ancestral to the selected US landraces through northern Mexican and southwestern US landraces (Merrill et al., 2009; Sánchez, 2011). The US landraces selected for this study are early, parental landraces (Northern Flint and Southern Dent) used to create the early, US Corn Belt inbreds and hybrids (Troyer, 1999; Labate et al., 2003; van Heerwaarden et al., 2012).

**Table 1.**
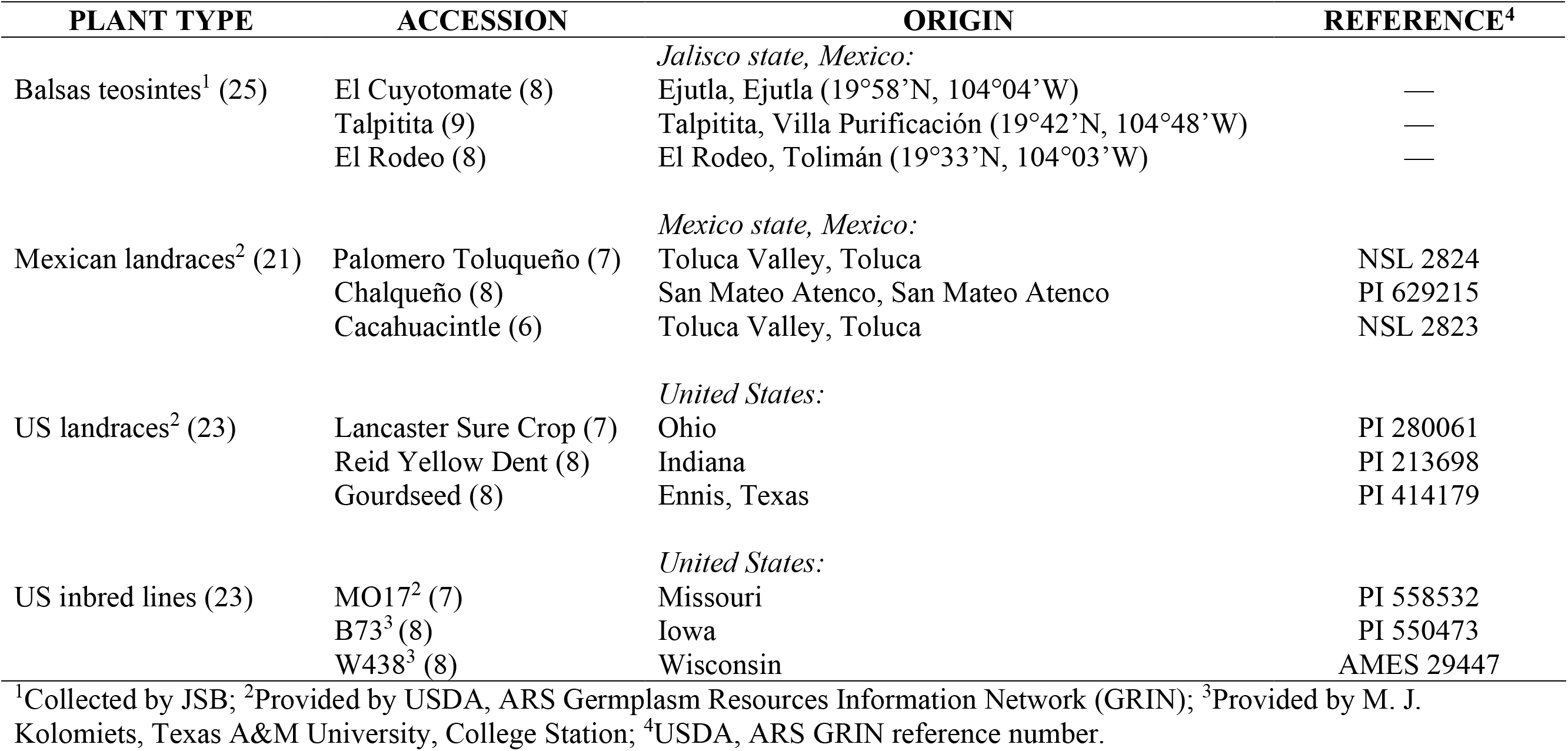
Plant types, accessions and number of bio-replicates per each (in parenthesis), and their geographic origins and reference numbers. From top to bottom, the plant types and their locations of origin span the domestication, spread, and breeding processes of maize from Mexico to the US Corn Belt.

Seeds of each accession were germinated in disposable Petri dishes (150×15mm) within moistened paper towels for 3 d. Teosinte seeds were initiated 1 d before maize seeds because they required more time to germinate, and were removed from their fruitcases with a nail clipper. Preliminary germination assays showed no need for seed surface sterilization. After germination, individual seedlings were transplanted to cone-tainers (4×25 cm diameter×length) (Stuewe & Sons, Tangent, OR, USA) and grown for additional 10-12 d; water was provided as needed. The cone-tainers were modified with chiffon mesh covering the bottom to prevent escape of Western corn rootworm larvae (preliminary assays not shown here). Growing conditions were 25 ± 2°C, 50% RH, and 12:12 photoperiod (L:D). The soil used was Baccto® premium potting soil (Michigan Peat Co., Houston, TX, USA), and was sifted (60 mesh strainer) to facilitate subsequent washing of roots (see below). The number of biological replicates per treatment (= seedlings) used for all assays were as follow: Balsas teosinte, n = 25, Mexican landraces, n = 21, US landraces, n = 23, and US inbred lines, n = 23.

WCR eggs (diapause strain) were provided by USDA-ARS-North Central Agricultural Research Laboratory (Brookings, SD, USA). Eggs were incubated in Petri dishes at 25 ± 2°C, ~ 80% RH for 12 ± 1 d over moistened absorbing paper. Neonate 1^st^-instar larvae (< 24 h after eclosion) were used in all assays.

### Host Plant Resistance and Tolerance Assays

#### Plant Resistance

The aim of this assay was to assess plant resistance through insect and plant performance variables, and compare between pairs of plant types representing the domestication, spread, and breeding transitions in maize. We expected to find decreasing resistance from Balsas teosinte to US inbred lines, manifested as both enhanced WCR larval performance and increased seedling growth.

To assess WCR performance, 10 neonate WCR larvae were placed in each cone-tainer holding a ~15 d-old seedling, and allowed to feed for 10 d (Robert et al., 2012b); each seedling was paired with a control seedling of similar size and equal number of leaves in order to estimate seedling growth ratios, as explained below. After 10 d, the cone-tainer soil was carefully examined and WCR larvae were recovered, counted and stored in 75% EtOH. Subsequently, each larva’s head capsule width was measured to record whether they were in their 1^st^, 2^nd^, or 3^rd^ instar (Hammack et al., 2003). These measurements were made with a dissecting stereoscope at 75× magnification, and equipped with an eyepiece reticle ruler with 100 subdivisions within 10 mm, which had been previously calibrated with a micrometer. Following these measurements, larvae from each cone-tainer were placed in a vial, dried to constant weight (≥ 2 days at 65 °C), and weighed to obtain average weight per larva per each cone-tainer. Each cone-tainer represented a replicated sample for a plant type.

To assess plant performance, true-leaves 2 and 3 (from the bottom, exclusive of cotyledon) were excised from each seedling, and scanned to measure their surface area using ImageJ® software (Rasband, 2017). After this, the seedling was cut at the base of its stem, placed in a paper envelope (together with the corresponding excised leaves) and dried to constant weight (≥ 2 days at 65 °C) (Becker and Meinke, 2008). Seedling roots were rinsed under running water while gently rubbed to remove soil particles, and also dried to constant weight. Stem diameter for each seedling was measured before infestation with WCR, and again prior to harvesting of seedlings, using a digital micrometer (Pittsburgh^®^, Harbor Freight Tools, Camarillo, CA, USA). These measurements were used to assess seedling growth rate and lost seedling growth under WCR herbivory, as explained below.

A multivariate analysis of variance (MANOVA) was applied to evaluate whether resistance differed among the four plant types, indicating effects of domestication, spread, and breeding. The independent variables were ‘plant type’ (Balsas teosintes, Mexican landraces, US landraces, US inbred lines), and ‘accessions’ (three per plant as described above in *Plants*) which were nested within plant type in the MANOVA model. The dependent variables were foliar weight (leaves and stem), leaf surface area, root weight, larval survivorship (number of recovered larvae/10 initial larvae), and average larval weight (per cone-tainer); additionally, growth rate (= the ratio between seedling stem diameter at days 0 and 10), and lost growth (= the ratio between seedling stem diameter of WCR-infested and -noninfested seedlings at day 10 of the assay) were estimated, and included in the analyses. These growth ratios were used to account for known differences in seedling size among plant types (Chinchilla-Ramírez et al., 2017). All data were transformed to ln(*x*) prior to analyses; prior to ln(*x*)-transformation, surface area data were converted to square-root values, and weight data to cubic-root values. *A priori* contrasts were used for paired comparisons between Balsas teosintes and Mexican landraces, Mexican landraces and US landraces, and US landraces and US inbred lines, using a Sidak-adjusted significance level of *P* < 0.017 (Abdi, 2007). Pearson correlations of canonical scores with dependent variables were used to determine the contributions of each dependent variable to the total variation in the canonical axes of MANOVA’s centroid plots; Pearson’s *r* values ≥ |0.50|, and *P* ≤ 0.05 were considered significant.

Analysis of variance (ANOVA) was performed for each dependent variable (*P* < 0.05), except for the frequencies of WCR larval instars per plant type. Ratios of plant dependent variables (WCR-infested/noninfested) were used to avoid bias due to phenotypic differences between plant types, as explained above. ANOVA was followed by *a priori* contrasts to compare between pairs of plant types, as described above. *G*-tests were performed (*P* ≤ 0.017, per Sidak’s correction) to test whether the frequency distributions of WCR larval instars varied between pairs of plant types (Abdi, 2007). Additionally, the proportions of 3^rd^-instar larvae were calculated for each plant type, and used as a proxy for WCR developmental speed; comparisons between plant types were made using *a priori* contrasts (*P* ≤ 0.017). All statistical analyses were performed using JMP software (SAS Institute Inc., 2018).

### Plant Tolerance

The aim of this assay was to compare plant tolerance between plant types by measuring plant growth in presence and absence of WCR larvae. As before, the comparisons between plant types sought to assess the effects of domestication, spread, and breeding, as described above for *Plant resistance*. We expected to find increasing tolerance from Balsas teosintes to US inbred lines, manifested as compensation for tissue loss due to feeding by WCR larvae.

The methodology used to assess plant tolerance followed that of an earlier study, with appropriate modifications (Chinchilla-Ramírez et al., 2017). The plant variables measured for plant resistance (foliar weight, leaf surface area, final stem diameter, and root weight; see above) were measured in treated (with 10 WCR larvae) and control (without WCR larvae) seedlings. Control seedlings were plants similar in size and number of leaves to treated seedlings, so that each treated seedling had a paired, control seedling. MANOVA and Pearson correlations of canonical scores were conducted as described above under *Plant Resistance*, with some exceptions. Independent variables included ‘plant type’ (Balsas teosintes, Mexican landraces, US landraces, US inbred lines), ‘herbivory’ (with and without WCR larvae), ‘accessions’ (three per plant as described above in plants) nested within plant type, initial stem diameter (at 0 days) (as covariate), and the interaction term ‘herbivory × plant type;’ initial stem diameter was included to account for anticipated size different across plant types and accessions (Chinchilla-Ramírez et al., 2017). The dependent variables included were final stem diameter (at 10 days of the assay), foliar weight, leaf surface area, and root weight. Following MANOVA, *a priori* contrasts between plant types were used to separate multivariate means between pairs of plant types (critical *P* < 0.017, per Sidak’s correction), as described above. To examine whether seedlings compensated tissue lost to herbivory by WCR, we calculated the mean ratios (= weight of infested seedlings/weight of non-infested seedlings) for each dependent variable, and applied one-sample *t*-tests with the null hypothesis that ratios would not differ from 1 (i.e. H_0_ = 1, no loss nor gain of tissue with WCR herbivory); the critical significance level was set to *P* < 0.012, per Sidak’s correction for four tests (Abdi, 2007). We considered ratio values < 1 as indicative of under-compensation, values = 1 of compensation, and > 1 of over-compensation. Data for these comparisons were transformed to cubic root(*x*) values for analyses. All statistical analyses were performed using JMP software (SAS Institute Inc., 2018).

### Plant resistance-Plant tolerance trade-off

To address the hypothesis that plant resistance trades off with plant tolerance (i.e. are negatively correlated) we conducted correlation analysis of data obtained in the *Plant Resistance* and *Plant Tolerance* assays described above. Specifically, we estimated the per-plant accession means for WCR larva weight from the *Plan resistance* assay, and the per-plant accession mean differences in foliar weight between infested (with WCR larvae) and control (without WCR larvae) seedlings in the *Plant tolerance* assay. We considered larva weight as a proxy for resistance, and the difference in foliar weight as a proxy for tolerance; the difference in foliar weight, rather than the difference in root weight, was used as a tolerance proxy to preclude the effect of lost root tissue due to WCR feeding on any gain of root tissue due to compensation. Mean larva weights were converted to cube-root(*x*) values, and differences in foliar weight to ln(*x*) values to comply with the expectation of normality. Our null hypothesis was that Pearson’s correlation coefficient, *r*, was larger than −0.5, i.e. *r* > [−0.5, 1] at *P* ≤ 0.05, indicating the absence of a negative correlation.

## RESULTS

### Plant Resistance

Through MANOVA we assessed whether insect and plant performances were affected by plant type (Figure 1). The analysis revealed a significant multivariate effect on both plant type (Wilks’ λ = 0.365, *P* < 0.001) and accession nested within plant type (λ = 0.361, *P* = 0.037). *A priori* contrasts between plant types showed significant differences between Balsas teosintes and Mexican landraces (F_7, 69_ = 4.489, *P* < 0.001) (i.e. a domestication effect) as well as for Mexican landraces and US landraces (F_7, 69_ = 2.643, *P* = 0.017) (i.e. a geographical spread effect), but not between US landraces and US inbred lines (F_7, 69_ = 1.894, *P* = 0.083) (i.e. a non-significant breeding effect). The vertical axis in the canonical plot explained 82% of the variation, with root (r = 0.814, *P* < 0.001) and foliar (r = 0.766, *P* < 0.001) weights as the variables that contributed the most to the separation between plant types, whereas the horizontal axis explained 12% of the variation between plant types, with foliar weight (r = 0.526, *P* < 0.001) and plant growth (r = 0.519, *P* < 0.001) as the variables separating plant types (Figure 1).

**Figure 1.**
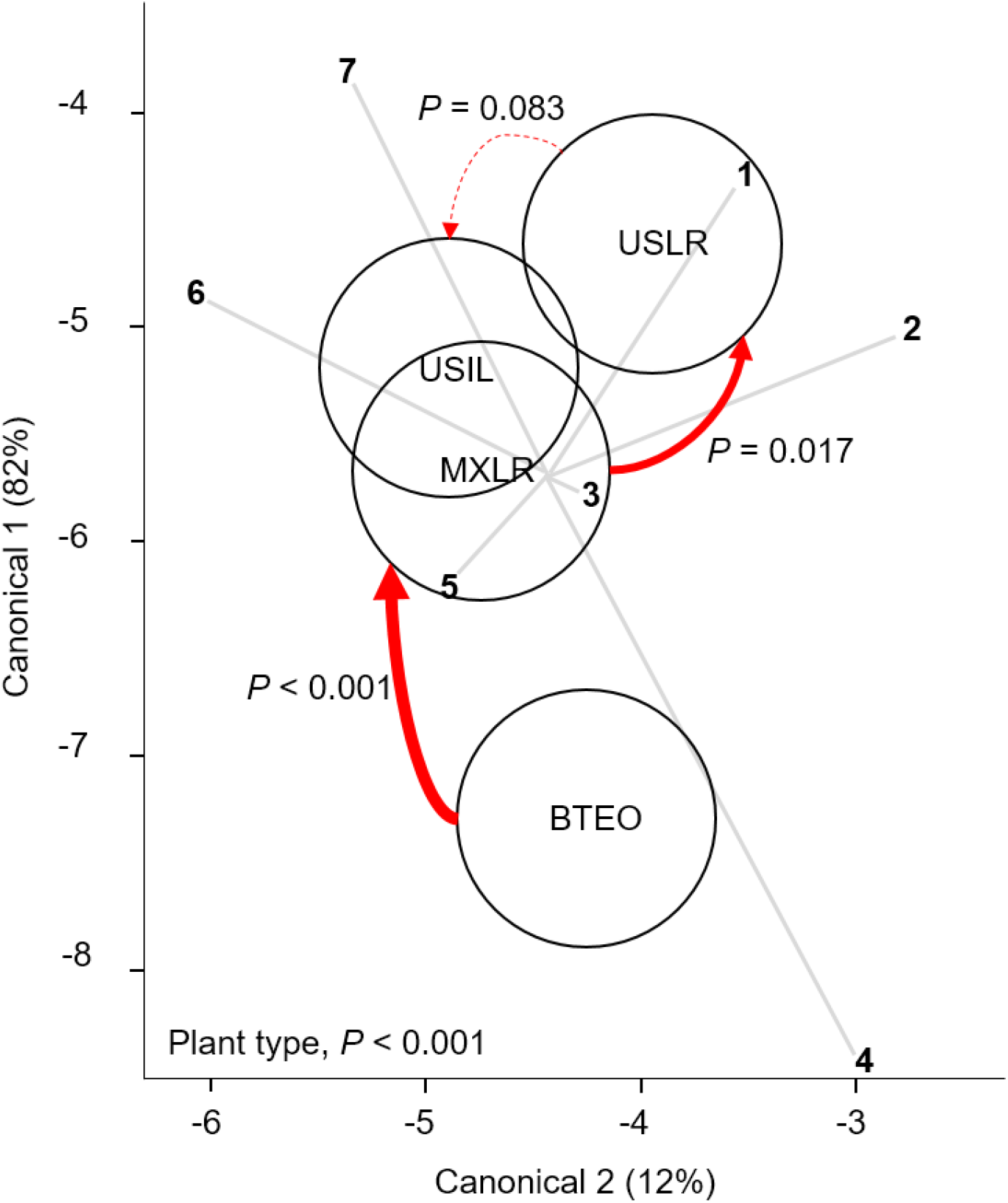
Canonical centroid plot from a Multivariate Analysis of Variance (MANOVA) (Wilks’ λ = 0.365, P < 0.001) for plant and Western corn rootworm variables associated with plant resistance; circles represent 95% confidence intervals around multivariate means for each plant type. The model included the independent variables ‘plant type’ (Balsas teosintes, Mexican maize landraces, US maize landraces, US inbred maize lines), and ‘accessions’ nested within plant type (three accessions per plant type, not shown here), and the dependent variables larval weight (ray 1), foliar weight (2), leaf surface area (3), plant growth (4), larval survivorship (5), root weight (6), and lost plant growth (7). Significant pair-wise comparisons between plant types (*a priori* contrasts with critical *P* of 0.017, per Sidak correction) are indicated by solid arrows (width is proportional to the confidence level); dashed arrow indicates a non-significant difference. The pair-wise comparisons are between plant types representing the domestication (BTEO vs. MXLR), spread (MXLR vs. USLR), and breeding (USLR vs. USIL) transitions evident in maize. BTEO = Balsas teosintes; MXLR = Mexican landraces; USLR = US landraces; USIL = US inbred lines.

Analysis of variance on each dependent variable revealed significant plant type effects on foliar ratio, root ratio, and larval weight, growth rate, and lost growth (*P* ≤ 0.026), but no effect on leaf surface area and larval survivorship (Table 2). *A priori* contrasts between plant types were applied to each significant dependent variable to assess domestication, spread, and breeding effects. These contrasts revealed significant differences between Balsas teosintes and Mexican landraces in foliar and root ratios (*P* ≤ 0.005); between Mexican landraces and US landraces in foliar ratio (*P* = 0.001), and; between US landraces and US inbred lines in foliar ratio, and larval weight (*P* ≤ 0.008) (Figure 2).

**Table 2.**
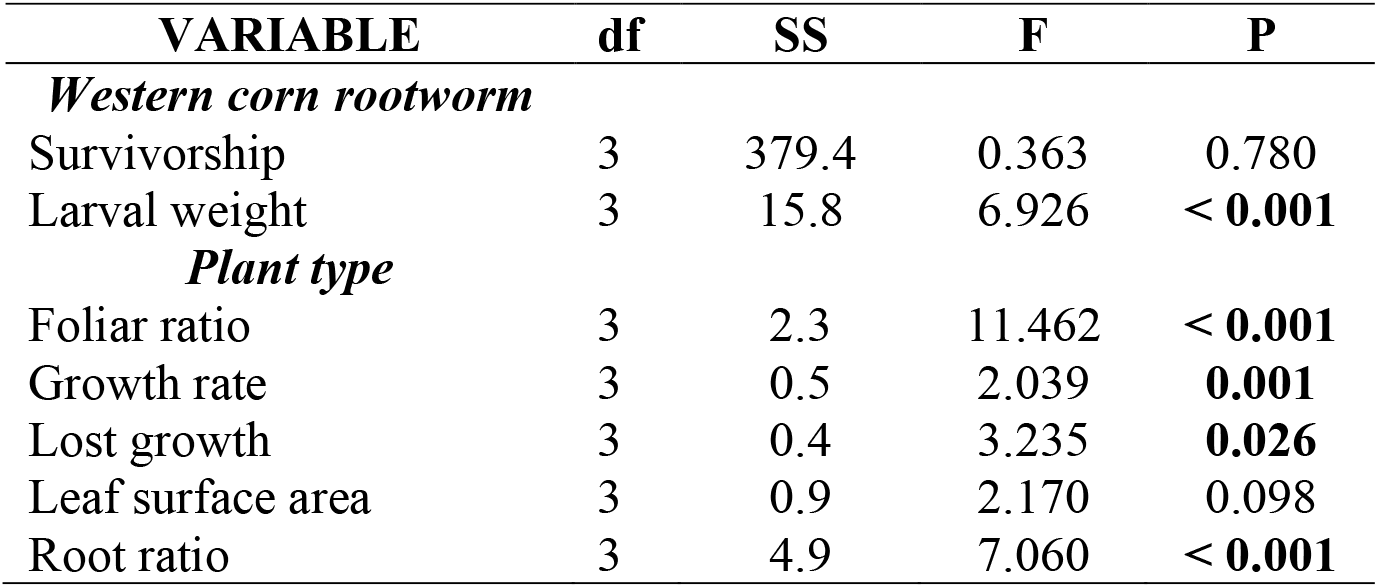
Analysis of variance (ANOVA) statistics for the independent variables ‘plant type’ (Balsas teosintes, Mexican maize landraces, US maize landraces, and US inbred maize lines) and seven plant and Western corn rootworm dependent variables associated with plant resistance. P values for variables significantly affected by plant type are shown in bold (P ≤ 0.05).

**Figure 2.**
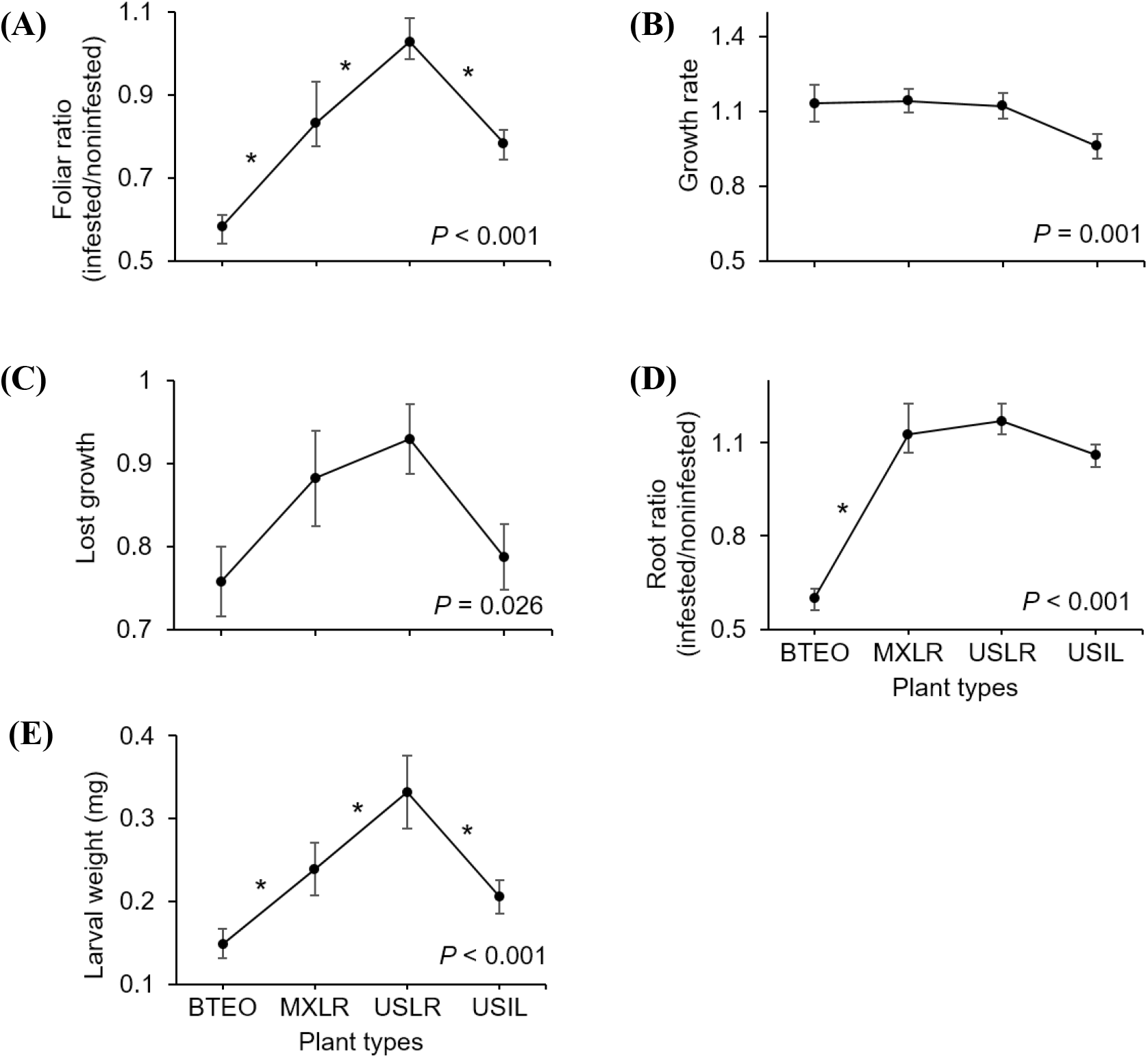
Paired comparisons between per-plant type means (± SE) of plant and Western corn rootworm (WCR) variables associated with plant resistance. Plant types are ordered left to right from most ancestral to most derived: Balsas teosintes (BTEO), Mexican maize landraces (MXLR), US maize landraces (USLR), and US maize inbred lines (USIL). Asterisks indicate significant difference (*a priori* contrasts with critical *P* ≤ 0.017, per Bonferroni correction) between means of contiguous plant types representing the domestication (BTEO vs. MXLR), spread (MXLR vs. USLR), and breeding (USLR vs. USIL) transitions in maize; univariate analysis of variance (ANOVA) *P* statistics are inset in each plot (see Table 2 for complete statistics). **(A)** Foliar ratio (= above-ground weights after 10 d, WCR-infested plants/noninfested plants); **(B)** Growth rate (= ratio between WCR infested seedling stem diameter at days 0 and 10 of the assay); **(C)** Lost growth (= stem diameter ratio after 10 d of WCR-infested plants/noninfested plants); **(D)** Root ratio (= belowground weights after 10 d of WCR-infested plants/noninfested plants). **(E)** Larval weight (= weights of WCR larvae after 10 d).

The distributions of larval instar frequencies varied among plant types (*G* = 40.43, 6 d.f., *P* < 0.001), (Figure 3A). Pairwise comparisons of frequency distributions showed significant differences between Balsas teosintes and Mexican landraces (*G* = 17.82, 2 d.f., *P* < 0.001), US landraces and US inbred lines (G = 17.32, 2 d.f., *P* < 0.001), but not between Mexican landraces and US landraces (*G* = 2.34, 2 d.f., *P* = 0.309), i.e. significant domestication and breeding effects, but not spread effects (Figure 3A). The development speed of WCR larvae did not differ significantly among plant types (F_3, 8_ = 2.33, *P* = 0.150) (Figure 3B).

**Figure 3.**
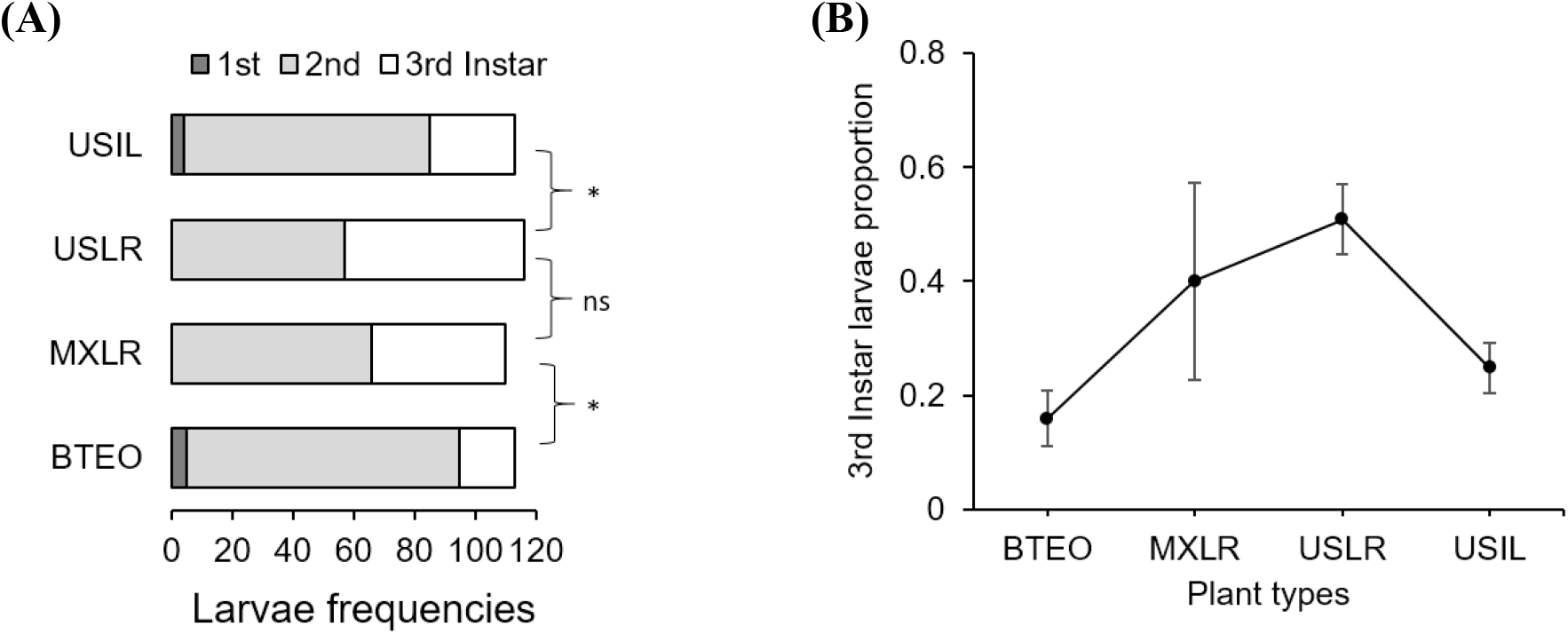
**(A)** Frequency distributions of 1^st^-, 2^nd^-, and 3^rd^-instar larvae, and **(B)** development speed (= proportion of larvae reaching 3^rd^-instar) of larvae of Western corn rootworm in trials concluding 10 d after neonates were allowed to feed on one of four plant types. Plant types were Balsas teosintes (BTEO), Mexican maize landraces (MXLR), US maize landraces (USLR), and US inbred maize lines (USIL), and are ordered from most ancestral to most derived. *A priori*, pair-wise comparisons between frequency distributions representing the domestication (BTEO vs. MXLR) and breeding (USLR vs. USIL) transitions were significant (* = G ≥ 17.25, *P* < 0.001), while the comparison representing the spread transition (MXLR vs. USLR) was not significant (G = 2.34, *P* = 0.309); the critical *P* value for these comparisons was set at P ≤ 0.017, per Sidak’s correction. Univariate analysis of variance (ANOVA) indicated that development speed did not vary across plant types (F_3, 8_ = 2.33, *P* = 0.150).

Overall, these results suggested that *Zea* resistance to WCR decreased with domestication and spread, and was partially recovered by breeding. Balsas teosintes appeared as the most resistant plant type, US landraces as the least resistant, and Mexican landraces and US inbred lines as intermediately resistant.

### Plant Tolerance

MANOVA (overall Wilk’s λ = 0.142, *P* < 0.001) revealed significant effects of herbivory (F_4, 156_ = 16.555, *P* < 0.001), plant type (λ = 0.622, *P* < 0.001), and herbivory × plant type interaction (λ = 0.869, *P* = 0.025) on seedling tolerance levels to WCR feeding (Figure 4). *A priori* contrasts within plant types revealed significant differences between WCR-infested and -noninfested Balsas teosinte (F_4, 164_ = 9.922, *P* < 0.001), Mexican landrace maize (F_4, 164_ = 4.115, *P* = 0.003), and US inbred maize (F_4, 164_ = 4.684, *P* = 0.001), but not within US landrace maize (F_4, 164_ = 2.253 *P* = 0.065) (Figure 4), suggesting that only US landraces displayed broad tolerance to WCR feeding. Correlation analysis of canonical scores showed that the vertical axis of the centroid plot explained 77% of the variation, with final stem diameter (r = 0.67, *P* < 0.001), foliar weight (r = 0.91, *P* < 0.001), and root weight (r = 0.90, *P* < 0.001) as the variables that most contributed to the separation between infested and non-infested plant types. The same analysis showed that the horizontal axis explained 19% of the variation between infested and noninfested plant types, with leaf surface area as the main explanatory variable (r = 0.53, *P* < 0.001) (Figure 4).

**Figure 4.**
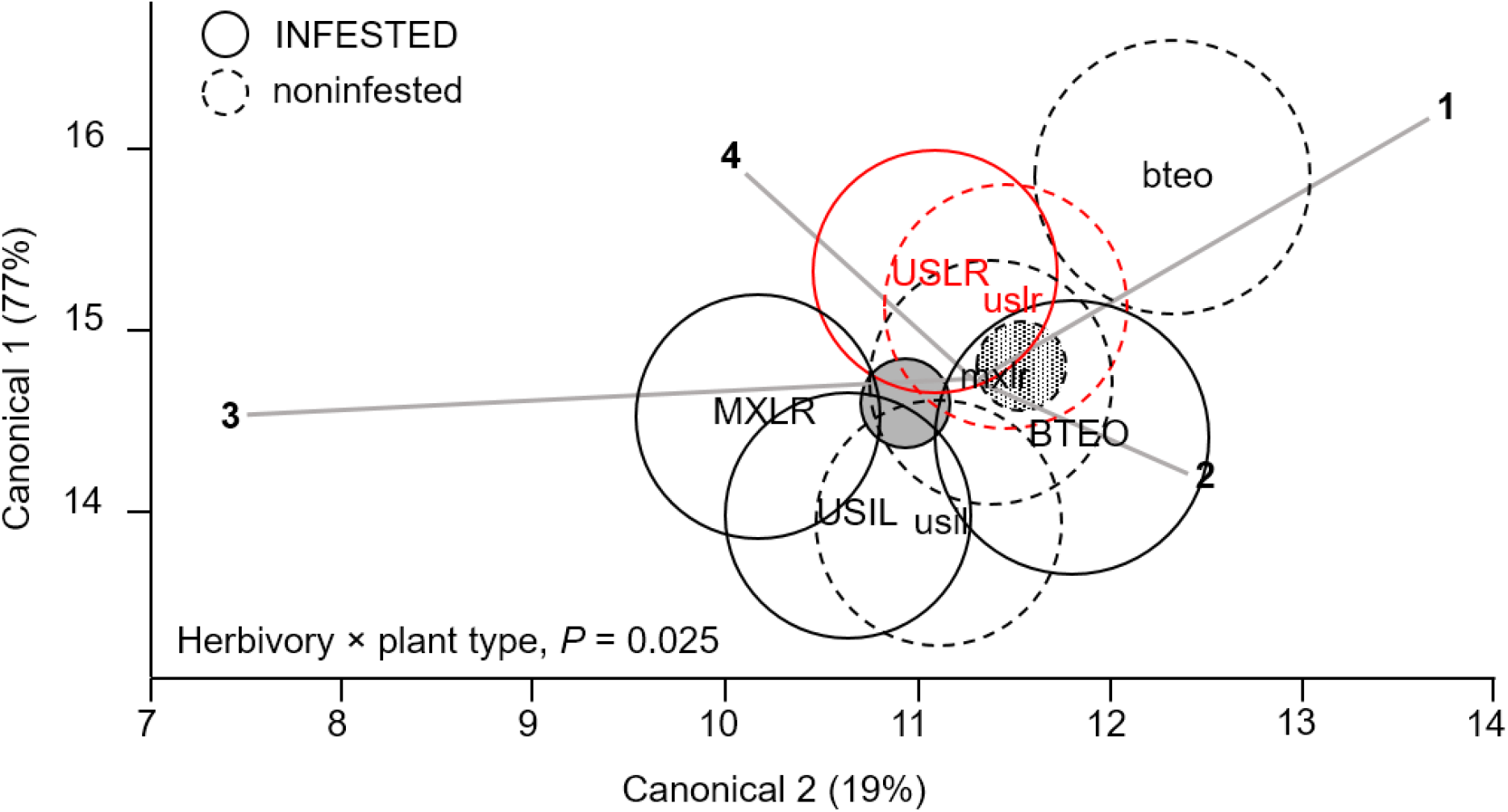
Canonical centroid plot for Multivariate Analysis of Variance (MANOVA) on plant variables associated with plant tolerance to feeding by Western corn rootworm; circles represent 95% confidence intervals around multivariate means for each plant type. The model includes the independent variables ‘plant type’ (Balsas teosintes, Mexican maize landraces, US maize landraces, US inbred maize lines), ‘accessions’ nested within plant type (three accessions per plant type, not shown here), herbivory (Western corn rootworm presence or absence) and the interaction term ‘herbivory × plant type.’ The dependent variables were foliar weight (ray 1), leaf surface area (2), final stem diameter (3), and root weight (4). The overall model (Wilks’ λ = 0.142, *P* < 0.001) and main effects were significant: plant type (λ = 0.622, *P* < 0.001), herbivory (F_4, 164_ = 16.555 *P* < 0.001), and herbivory × plant type (λ = 0.869, *P* = 0.025). Pair-wise comparisons between Western corn rootworm-infested and -noninfested plants within plant types (depicted by continuous circles/upper-text and dashed circles/lower-case text) were significant for all plant types, except for US landraces (red circles): Balsas teosintes, F_4, 164_ = 9.922, *P* < 0.001; Mexican landraces, F_4, 164_ = 4.115, *P* = 0.003; US landraces. F_4, 164_ = 2.253, *P* = 0.065; US inbred lines, F_4, 164_ = 4.684, *P* = 0.001. Smallest circles (filled) near plot center represent overall Western corn rootworm-infested (solid line and filling) and -noninfested (dashed line, patterned filling) plants. BTEO, bteo = Balsas teosintes infested or noninfested, respectively, by Western corn rootworm; MXLR, mxlr = Mexican landraces; USLR, uslr = US landraces; USIL, usil = US inbred lines.

Within each plant type, tissue losses, assessed as mean ratios (= WCR-infested seedlings/non-infested seedlings) of foliar weight, leaf surface area, final stem diameter, and root weight, were found to be undercompensated (i.e. ratio < 1.0, *P* < 0.001) in both Balsas teosintes and US inbred lines, with the exception of root tissue, which was compensated in US inbred lines (i.e. ratio > 1.0, *P* = 0.780) (Figure 5). Mexican landraces compensated foliar, final stem diameter and root tissue losses (i.e. ratio did not differ from 1.0, *P* ≥ 0.019), and undercompensated leaf surface area losses (P < 0.001). Finally, US landraces compensated all tissue losses, foliar, leaf surface area, final stem diameter, and root tissue (*P* ≥ 0.020). These results suggested that US landraces displayed tolerance to WCR as they consistently compensated tissue losses, Mexican landraces and US inbreds displayed partial tolerance, and Balsas teosintes did not display tolerance (Figure 5).

**Figure 5.**
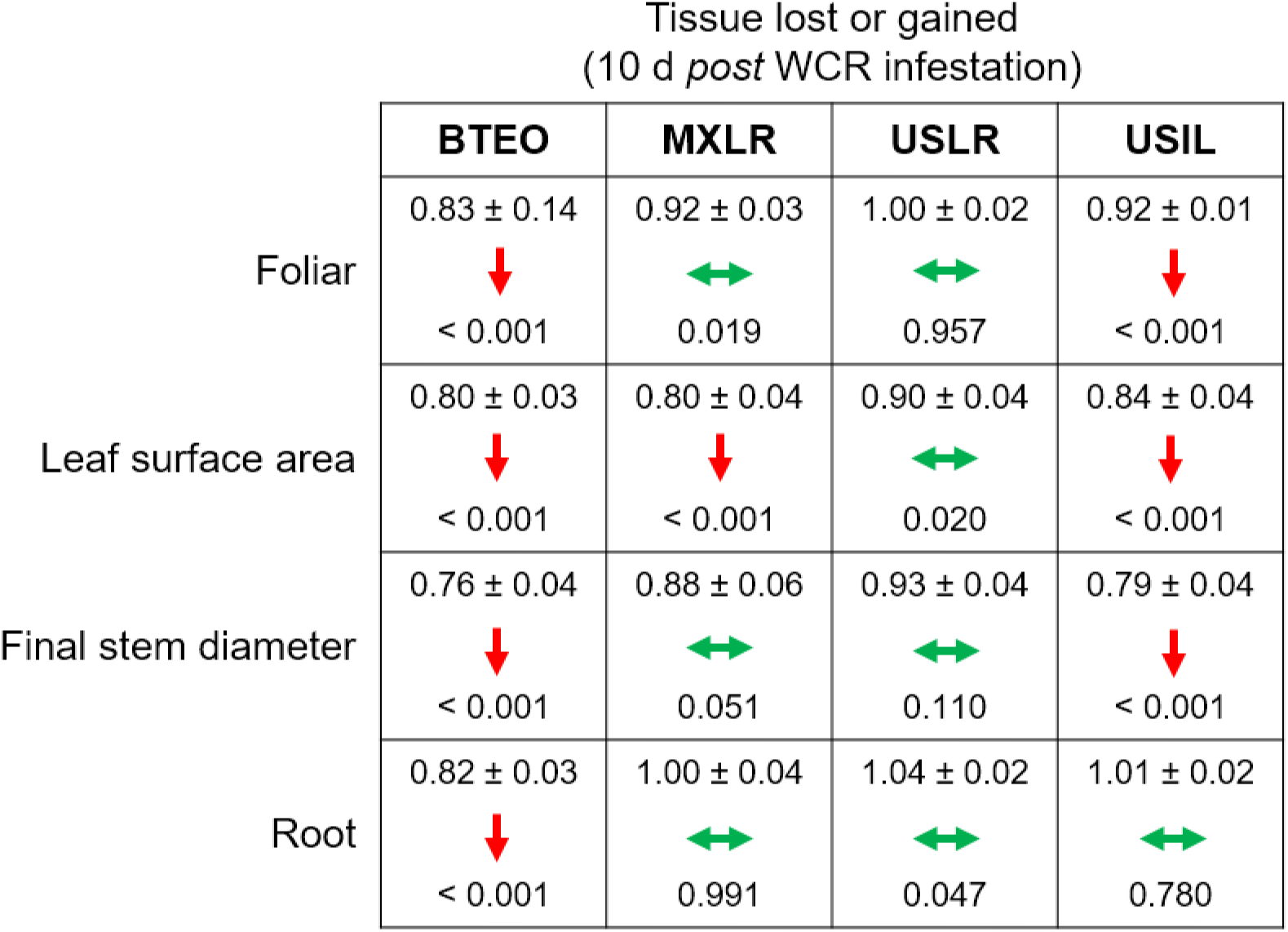
Effects of herbivory by Western corn rootworm on plant tolerance variables in four plant types, ordered from most ancestral to most derived: Balsas teosintes (BTEO), Mexican maize landraces (MXLR), US maize landraces (USLR), and US inbred maize lines (USIL). The plant tolerance variables are mean ratios (= Western corn rootworm-infested plants/noninfested plants) of final stem diameters, foliar weights, leaf surface areas, and root weights. One-sample *t*-tests were used to compare mean ratios for each plant type against expected ratio of 1.0, which indicates tissue compensation (i.e. no tissue lost or gained in Western corn rootworm-infested plants relative to noninfested plants); the mean ratio (± SE) and *P* value are shown within each cell. Within each cell, double-pointed, horizontal green arrows indicate compensation (mean ratio does not differ from 1.0), and downward, red arrows indicate undercompensation (mean ratio < 1.0). Critical *P* for each *t*-test was set at *P* ≤ 0.012, per Sidak’s correction.

Herbivory × plant type interaction effects are shown in Figure 6. Significant differences between infested and noninfested seedlings were found for foliar (F_3, 167_ = 3.126, *P* = 0.027) and root (F_3, 167_ = 4.039, *P* = 0.008) weights, but not for final stem diameter (F_3, 167_ = 0.8140, *P* = 0.487) nor leaf surface area (F_3, 167_ = 0.471, *P* = 0.702). *A priori* contrast comparisons between infested and noninfested seedlings (*P* ≤ 0.012; Sidak corrected) revealed significant foliar tissue losses (i.e. undercompensation) in Balsas teosintes (F_1, 167_ = 27.536, *P* < 0.001) and Mexican landraces (F_1, 167_ = 7.543, *P* = 0.007), while US landraces (F_1, 167_ = 0.890, *P* = 0.346) and US inbred lines (F_1, 167_ = 4.127, *P* = 0.041) did not lose nor gain tissue (i.e. compensation) (Figure 6a). *A priori* contrast comparisons for root weights revealed that Balsas teosintes lost tissue (i.e. undercompensation) (F_1, 167_ = 13.576, *P* < 0.001), whereas Mexican landraces (F_1, 167_ = 0.005, *P* = 0.942), US landraces (F_1, 167_ = 0.424, *P* = 0.515), and US inbred lines (F_1, 167_ = 0.158, *P* = 0.691) did not lose nor gain tissue (i.e. compensation) (Figure 6b).

**Figure 6.**
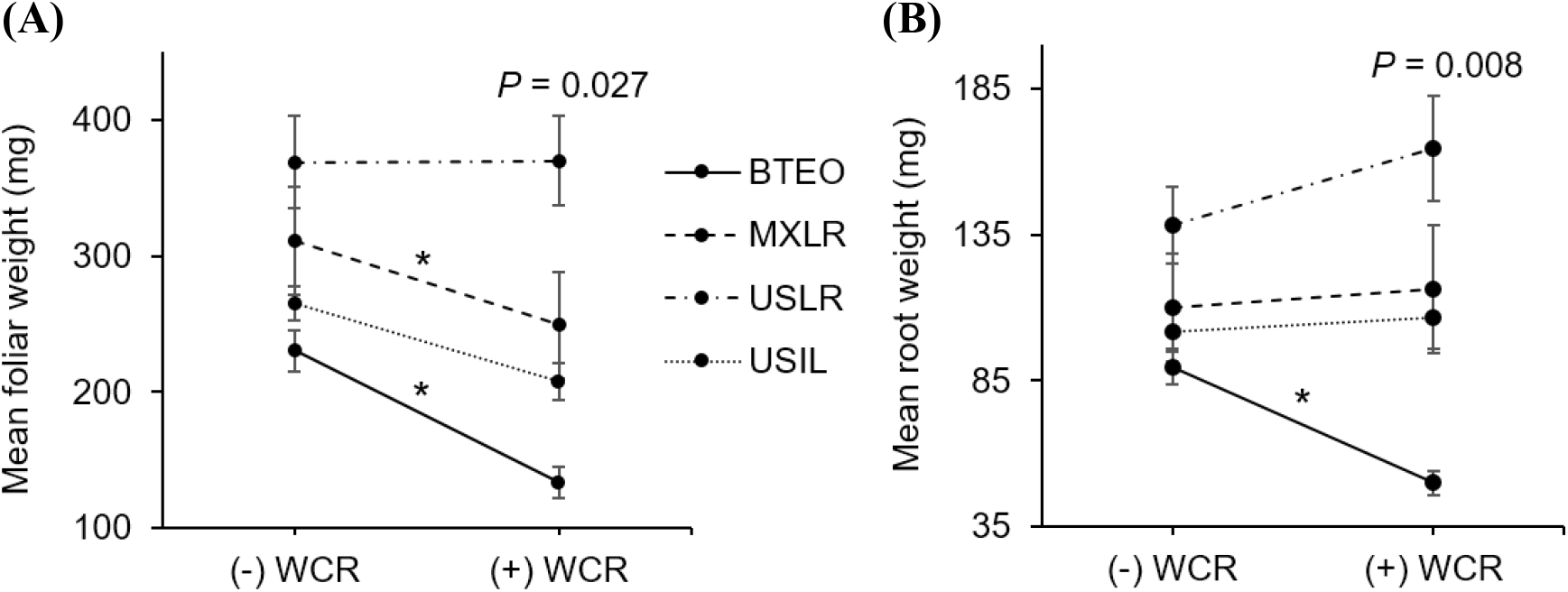
**(A)** Above- and **(B)** belowground tissue losses in four plant types, Balsas teosintes (BTEO), Mexican maize landraces (MXLR), US maize landraces (USLR), and US inbred maize lines (USIL), exposed to root herbivory by Western corn rootworm (WCR) larvae. Inset in each plot are the univariate analysis of variance (ANOVA) statistics for the herbivory (+WCR, -WCR) × plant type effect in foliar weight (F_3, 167_ = 3.126, *P* = 0.027) and root weight (F_3, 167_ = 4.039, *P* = 0.008). Comparisons between plant types exposed (+WCR) and unexposed (-WCR) to Western corn rootworm larvae were made via *a priori* contrasts, with a critical *P* value for each paired comparison set at *P* ≤ 0.012, per Sidak’s correction. Significant herbivory effects are indicated by an asterisk (*).

Overall, these results suggested that *Zea* tolerance to WCR was gained with domestication and reinforced by spread. However, it also suggested that breeding weakened tolerance to a point comparable to that evident in Mexican landraces. The tolerance levels, ordered from most to least tolerant plant type appeared to be US landraces, Mexican landraces, and US inbreds, while Balsas teosintes appeared to be intolerant.

### Plant resistance-Plant tolerance trade-off

Correlation analysis showed a significant negative correlation between per-plant accession larval weights and differences in foliar weights between WCR-infested and non-infested seedlings (r = −0.646, *P* = 0.023) (Figure 7). Consistent with the *Plant resistance* and *Plant tolerance* results, the analysis suggested that Balsas teosintes was the most resistant plant type, US landraces was the least resistant, and Mexican landraces and US Inbred lines were intermediately resistant. Conversely, it suggested that US landraces was the most tolerant plant type, Balsas teosintes was the least tolerant, and Mexican landraces and US Inbred lines were intermediately tolerant. Overall, these results suggested that resistance declines with increasing tolerance in *Zea*.

**Figure 7.**
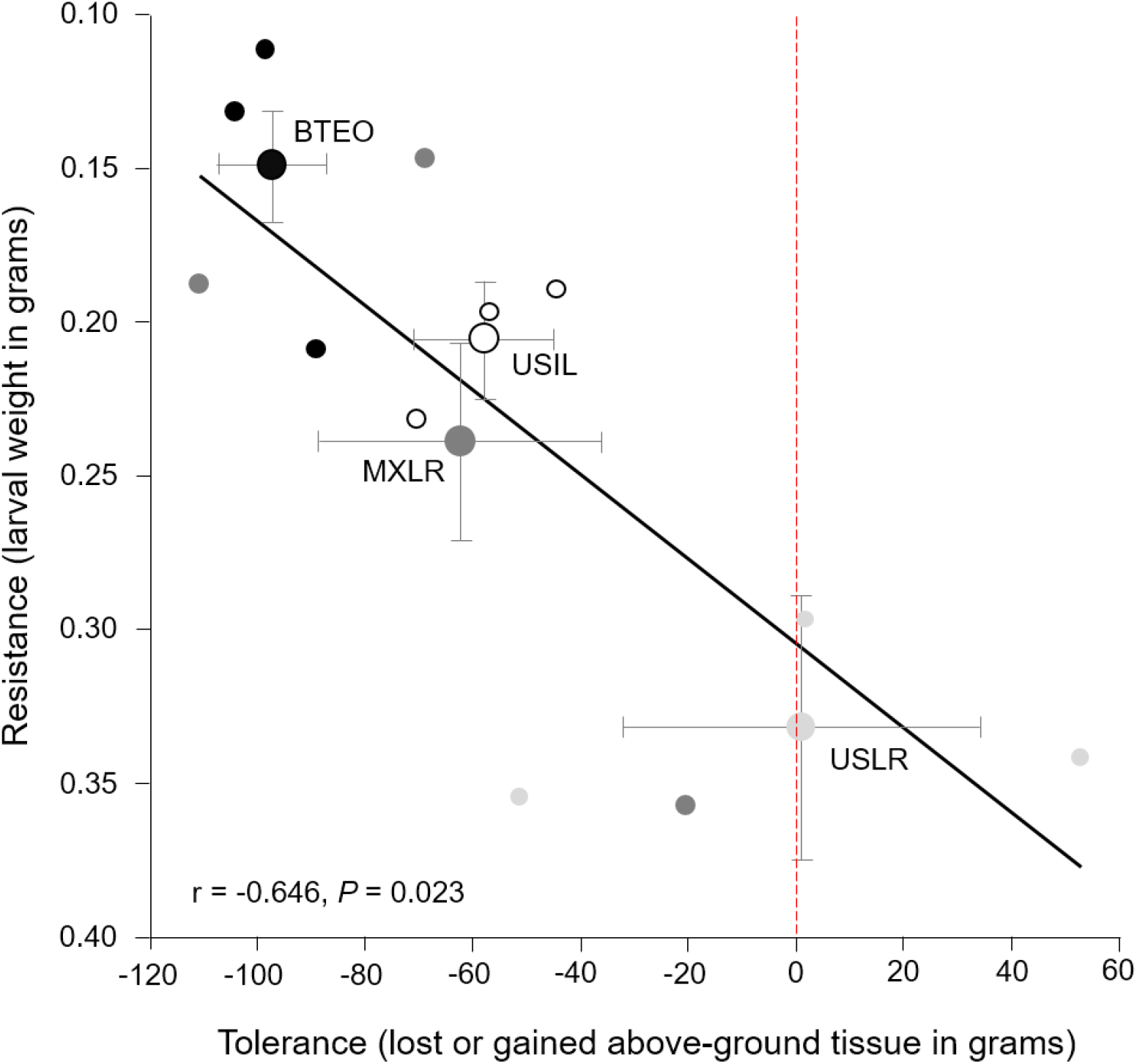
Relationship between resistance (expressed as larval weight) and tolerance (expressed as plant tissue loss or gain) to root herbivory by Western corn rootworm larvae in 12 plant accessions (small circles), with three accessions corresponding to each of four plant types (large circles with bi-directional standard error bars). Note that Y-axis values increase from top to bottom. Plant types are Balsas teosintes (BTEO), Mexican maize landraces (MXLR), US maize landraces (USLR), and US inbred maize lines (USIL). The weight of Western rootworm larvae (g) after 10 days of feeding on each accession was used as a proxy for resistance, while the loss or gain of above-ground tissue (g) of each accession after 10 days of exposure to root herbivory by rootworm larvae was used as a proxy for tolerance. Inset is Pearson’s correlation *r* statistic corresponding to the 12 plant accessions. The red, dotted vertical line on *x*-axis indicates tissue compensation (i.e. no tissue lost nor gained); means to the left of the dotted line are suggestive of undercompensation for tissue loss, and means to the right are suggestive of overcompensation for tissue loss.

## DISCUSION

This study addressed whether maize defense, in the forms of resistance and tolerance to root herbivory, was mediated by domestication, spread, and breeding processes that spanned divergent environments and thousands of years and kilometers. To that end, we studied resistance and tolerance to Western corn rootworm feeding in four host plants that encompass those processes: Balsas teosinte, Mexican landrace maize, US landrace maize, and US inbred line maize. Specifically, we assessed the performances of WCR larvae and host plant types as proxies for resistance, and the performances of host plant types as affected by WCR feeding as proxies for tolerance. We expected to find that maize resistance against WCR was weakened with domestication, spread, and breeding, and that tolerance to WCR increased as resistance decreased. Our results were consistent with our expectations, though not entirely. On one hand, maize resistance indeed decreased from Balsas teosintes to US landraces, i.e. with maize domestication and spread, though, surprisingly, the trend seems to have reversed with breeding: US inbred lines showed more resistance to WCR than their US landrace ancestors, so were intermediately resistant rather than least resistant. On the other hand, tolerance indeed increased as resistance decreased, as expected.

### Maize resistance decreased with domestication and spread, buy increased with breeding

Our results suggested that maize resistance to root herbivory by WCR was weakened with domestication and spread, as we expected, while breeding affected resistance differently than we expected. Specifically, MANOVA revealed a strong multivariate effect of plant type on resistance variables, and *a priori* comparisons showed significant differences between Balsas teosintes and Mexican landraces, as well as between Mexican landraces and US landraces, but not between US landraces and US inbreds. Similarly, ANOVAs of individual dependent variables showed both domestication and spread effects, especially on WCR larval performance (i.e. weight), which was enhanced on Mexican landraces compared to Balsas teosintes, as well as on US landraces compared to Mexican landraces. However, WCR larval performance declined on US inbreds compared to US landraces, in partial contrast to our MANOVA results. Moreover, an *a posteriori* contrast comparison between between Balsas teosintes and US inbred lines showed no significant differences in larval weight and lost plant growth (F_1, 167_ = 4.033, *P* = 0.046; data not shown; Sidak-corrected critical *P* ≤ 0.012). This result may indicate significant allocation of resources to defense against WCR in US inbred lines, as would be expected to support enhanced resistance. Overall, these results suggested that domestication and spread affected resistance, as we anticipated and in agreement with other studies (Bazzaz et al., 1987; Rosenthal and Dirzo, 1997; Rodriguez-Saona et al., 2011), but resistance was partly recovered with breeding, contrary to expected. The optimal plant defense hypothesis predicts that there is a cost of defense, particularly that metabolic resources cannot be simultaneously used to defend, grow, and reproduce, so that plant fitness increases when herbivory decreases or is absent (Stamp, 2003). This prediction did not seem to apply to US inbred lines, which appeared to compensate root tissue (see below) while decreasing WCR larval weight.

Domestication, spread, and breeding significantly affected WCR performance. These results suggested two, non-exclusive defense strategies related to plant defense biochemistry. First, the nutritional value for WCR in *Zea* host plants may have increased from Balsas teosinte to US landrace maize, but decreased in US inbred maize. Changes in nutrient composition may cause differences in larval weight and development, while maintaining survivorship (Meihls et al., 2018). WCR uses a blend of sugars and fatty acids, but not amino acids, as phagostimulants to accept and feed on maize (Bernklau and Bjostad, 2008). Sucrose, although of non-nutritional value to most insects, is known to be an important phagostimulant, including for larvae of Coleoptera, and may be more relevant for host plant acceptance or rejection than any amino acid considered important for insect development (Chapman, 2003). There are no direct studies, to our knowledge, comparing root nutritional value among *Zea* plants. However, *Zea* has experienced selection in 2-4% of its genome, resulting in numerous biochemical differences among teosintes, landraces, and inbred lines (Dorweiler et al., 1993; Wright et al., 2005; Flint-Garcia et al., 2009; de Lange et al., 2014). Secondly, maize landraces may be down-regulating some secondary metabolites, while maize inbreds may be up-regulating them to levels similar to those in Balsas teosinte. The composition of secondary metabolites has been altered by domestication in various crops, affecting their interactions with specialist and generalist insects (Da Costa and Jones, 1971; Howe et al., 1976; Gols et al., 2008; Chacon-Fuentes et al., 2015). Typically, generalist herbivores perform better on domesticated plants compared to their wild relatives due to a reduction in secondary metabolites (Rodriguez-Saona et al., 2011; Bellota et al., 2013; Szczepaniec et al., 2013; Turcotte et al., 2014; Bernal et al., 2015; Chen et al., 2015b; Maag et al., 2015). WCR shifted to maize when the crop reached northern Mexico (Lombaert et al., 2017), and encountering maize landraces with weaker defenses than its original, wild host may have been advantageous for the quasi-specialist WCR (Branson and Ortman, 1967; 1970; Hahn and Maron, 2016). Maize breeding, conversely, may have partly reversed the decreasing trend of secondary metabolite levels, without a concurrent effect on maize productivity. Maize per-plant productivity (but not per-area yields) seems to have reached a maximum several decades ago, so that any productivity costs of chemical defense may be negligible, while concurrent breeding efforts may have inadvertently selected for WCR resistance, as evident for other maize pests (Duvick, 2005). Regardless of the relative importance of either defense strategy, the differences in WCR and seedling responses among plant types in our study was consistent with hypotheses of resistance reductions with domestication and spread (Rosenthal and Dirzo, 1997; Whitehead et al., 2017; Zust and Agrawal, 2017). However, breeding seemingly increased resistance (measured as decreased WCR performance) in US inbred lines, with no apparent cost to productivity. Further below, we discuss conditions under which resistance against WCR may have increased in US inbred maize concurrently with productivity, i.e. yield gains, particularly in the context of intensive maize agriculture reliant on modern technologies, such as synthetic fertilizers and pesticides, among others.

### Maize tolerance increased with domestication and spread, but decreased with breeding

Our results suggested that maize tolerance of root herbivory by WCR was enhanced as resistance decreased with domestication and spread, as expected, while breeding affected tolerance (and resistance) differently than expected. Specifically, MANOVA revealed strong multivariate effects of plant type on tolerance variables, and contrast comparisons revealed increasingly smaller (but significant) differences between WCR-infested and control seedlings (as indicated by *F* and *P* values) in Balsas teosintes, Mexican landraces, and US inbreds, while a significant difference was not found in US landraces. In this regard, US landraces showed the smallest partial *η*^2^ effect size of WCR infestation on seedling growth (partial *η*^2^ = 0.055, 0.000 – 0.101), while effect sizes were 3.6- (partial *η*^2^ = 0.200, 0.099 – 0.273), 2.4- (partial *η*^2^ = 0.134, 0.046 – 0.200), and 1.9-fold greater (partial *η*^2^ = 0.107, 0.027 – 0.168) in Balsas teosintes, Mexican landraces, and US inbreds, respectively (data not shown) (Richardson 2011). Similarly, our univariate analyses showed that US landraces consistently compensated for tissue losses, while Mexican landraces and US inbreds inconsistently compensated for tissue losses, and Balsas teosintes consistently undercompensated for tissue losses. Finally, measured as total above- or belowground tissue losses, Balsas teosintes lost both above- and belowground tissue with WCR feeding, Mexican landraces and US inbreds lost aboveground tissue, and US landraces compensated for above- and belowground tissue losses. Taken together, these results suggested that tolerance was strongest in US landraces, weakest in Balsas teosintes, and intermediate in Mexican landraces and US inbreds.

Domestication and subsequent farming could favor tolerance evolution when abiotic factors (e.g., soil nutrients, light availability) mediate the selection of plant defenses against herbivores (Hahn and Maron 2016). Annual crops, grown as they typically are, in resource rich environments are predicted to maximize fitness by allocating resources towards growth and reproduction, and trading-off constitutive resistance to herbivory (Herms and Mattson, 1992; Rosenthal and Dirzo, 1997). Additionally, biotic factors may impose selective pressures on domesticated plants. For example, in Hahn and Maron’s (2016) framework for intraspecific variation of plant defenses, two factors mediate defense evolution to tolerance or resistance: Low physiological stress (selecting for fast growing plants) and herbivory pressure (selecting for induced resistance). Moreover, herbivory pressure may indirectly select for tolerance as some root herbivores are able to manipulate the host to allocate primary metabolites (e.g., carbon, phosphorus, among others) to roots, and increase their host’s quality (Robert et al., 2012a). Such allocation may increase the likelihood of root compensation and, therefore, tolerance to root herbivory, and plants able to compensate for root herbivory may be favored by selection (cf. Figure 5 and 6). In parallel, this may explain the increased resistance and weakened tolerance in US inbred lines compared to US landraces. US inbred lines have been bred in contexts of low physiological stress and high WCR herbivory, especially since the 1940s, compared to the contexts in which their ancestral landraces were grown and selected (see below).

### Maize resistance and tolerance trade-off

Overall, our results showed a negative correlation between resistance and tolerance, consistent with optimal defense hypotheses and our expectation (Herms and Mattson, 1992; Blossey and Notzold, 1995; Zou et al., 2007; Hahn and Maron, 2016). However, we expected that this correlation would be consistent also with the evolutionary transitions between Balsas teosintes and US inbred lines. Usually, trade-offs are observed when fitness is compromised due to competing resource demands, e.g., resistance and fast growth (Agrawal et al., 2010). Natural selection may benefit one or the other depending on their direct or ecological costs (Strauss et al., 2002). Artificial selection, however, may favor a trade-off between a desired trait and a less-desired trait, e.g., selection for productivity weakened resistance, as our results suggested for Mexican landraces. A changing environment and herbivory pressure for US landraces may have led to an adaptive, negative correlation, where maize exposed to WCR under higher resource availability was subjected to strong selection for herbivory tolerance (Agrawal et al., 2010). Furthermore, plant resistance may de-escalate when a plant’s herbivore fauna is dominated by mono- or oligophagous insects, such as WCR (Agrawal and Fishbein, 2008; Agrawal et al., 2010). WCR became a pest after maize agriculture spread to North America, and may have been an important selection force shaping the defenses of modern maize in the US. The extended, thousands of years-association between maize and WCR — punctuated with severe WCR bottlenecks when maize agriculture became dominant in (current) southwestern (ca. 500 YBP) and northern (ca. 180 YBP) USA states — may have led to an evolutionary compromise, with maize gaining tolerance and WCR becoming a specialist (Robert et al., 2012a; Lombaert et al., 2017; Robert et al., 2017).

### Disarmed by agricultural intensification: Maize traded Western corn rootworm tolerance for token resistance

Our results addressing the effects of maize domestication and spread on defense strategy evolution were consistent with theoretical predictions concerning resistance and tolerance evolution in the contexts of plant productivity-resistance trade-offs and plant resistance-tolerance trade-offs, respectively (Agrawal et al., 2010; Pearse et al., 2017; Zust and Agrawal, 2017). Namely, our results showed that resistance to WCR decreased with both maize domestication and spread, and tolerance increased as resistance decreased, as expected. However, our results addressing the effects of breeding on maize defenses were inconsistent with predictions based on productivity-resistance and resistance-tolerance trade-offs. Specifically, breeding reversed the preceding trend of decreasing resistance and increasing tolerance so that US inbred lines were less tolerant and more resistant to WCR than their ancestral US landraces. We believe that this reversal is a result of agricultural intensification of maize production, particularly the systematic breeding of maize varieties for maximum yield under the umbrella of commercial, synthetic fertilizers, irrigation, and pesticides (Bernal and Medina, 2018). Under such intensification, pesticides provided relief from WCR injury without a metabolic cost to the crop, and fertilizers coupled with irrigation enhanced plant nutrient levels to support on one hand the productivity increases gained with systematic breeding, and on the other to offset any productivity losses due to WCR and other pests. This intensification period began in the late 1940s with the widespread availability of hybrid maize varieties, chemical fertilizers, and pesticides, and in the context of increasing pressure by WCR, which up to then had not been a significant pest (Perkins, 1982; Palladino and Fitzgerald, 1996; Duvick, 2005; Gray et al., 2009; Smith, 2011; Lombaert et al., 2017). In contrast, the period prior to intensification was characterized by natural and farmer (artificial) selection of maize landraces for broad resistance to environmental stresses, the absence of pesticides and commercial fertilizers, and minimal WCR pressure; this period ended with the deployment of commercial hybrid varieties, and decline of landraces, beginning in the 1930s (Duvick, 2005; Kutka, 2011; Smith, 2011; Bernal and Medina, 2018).

Overall, our results were consistent not only with predictions concerning plant defense evolution in the contexts of plant productivity-resistance trade-offs and plant resistance-tolerance trade-offs, as noted above, but also with predictions concerning defense strategy evolution in the context of variable resource availability and environmental stresses, particularly physiological stress (under low resource availability) and herbivory pressure (Herms and Mattson, 1992; Blossey and Notzold, 1995; Zou et al., 2007; Hahn and Maron, 2016) (Figure 8). We believe that shifts in resource availability, WCR pressure, and farmer selection of maize landraces to systematic breeding of maize inbred lines between the pre-intensification and intensification periods of maize production mediated the evolution of WCR defenses in US inbred maize lines (Duvick, 2005; Gray et al., 2009; Ivezić et al., 2009; Kutka, 2011; Smith, 2011; Lombaert et al., 2017; Mesa et al., 2017). For example, while the slight gain in WCR resistance evident in US inbred lines was not anticipated per expectations under a productivity-resistance trade-off, it was an anticipated result of directed systematic breeding for WCR resistance (and inadvertent selection under WCR pressure), and was associated with a loss of tolerance, as anticipated under a resistance-tolerance trade-off (Duvick, 2005; Agrawal, 2006; Agrawal and Fishbein, 2008; Gray et al., 2009; Ivezić et al., 2009; Agrawal et al., 2010). In Figure 8a, we show how resource availability may have increased (indicated by the arrow’s increasingly dark coloration) with maize domestication and spread, as maize — by that time an important food crop — is subjected to site selection and cultural practices aimed at enhancing its productivity. Concomitantly, physiological stress gradually may have lost importance as a driver of herbivore defense evolution as resource availability increased (see horizontal arrow at top of Figure 8, showing how resource availability is relevant to defense evolution at low resource availability, while herbivory pressure is relevant at high resource availability). In Figure 8B, we show how resource availability may have continued to increase and reached its highest level with the breeding transition, particularly with the advent of commercial fertilizers to support cultivation of high-yielding maize cultivars, i.e. intensification. At the same time, WCR emerged as an important pest of maize, and while it may have become a significant driver of herbivore defense evolution, its significance was mediated by the use of insecticides, which became widely available as maize agriculture was increasingly intensified. Altogether, we believe that our results illustrate how the evolution of defense strategies in maize, and perhaps other crops, is predicted by ecological-evolutionary hypotheses predicting defense strategy evolution in the contexts of plant resistance-productivity trade-offs, plant tolerance-resistance trade-offs, and varying resource availability vis-à-vis plant physiological stress and herbivory pressure (Herms and Mattson, 1992; Blossey and Notzold, 1995; Zou et al., 2007; Hahn and Maron, 2016).

**Figure 8.**
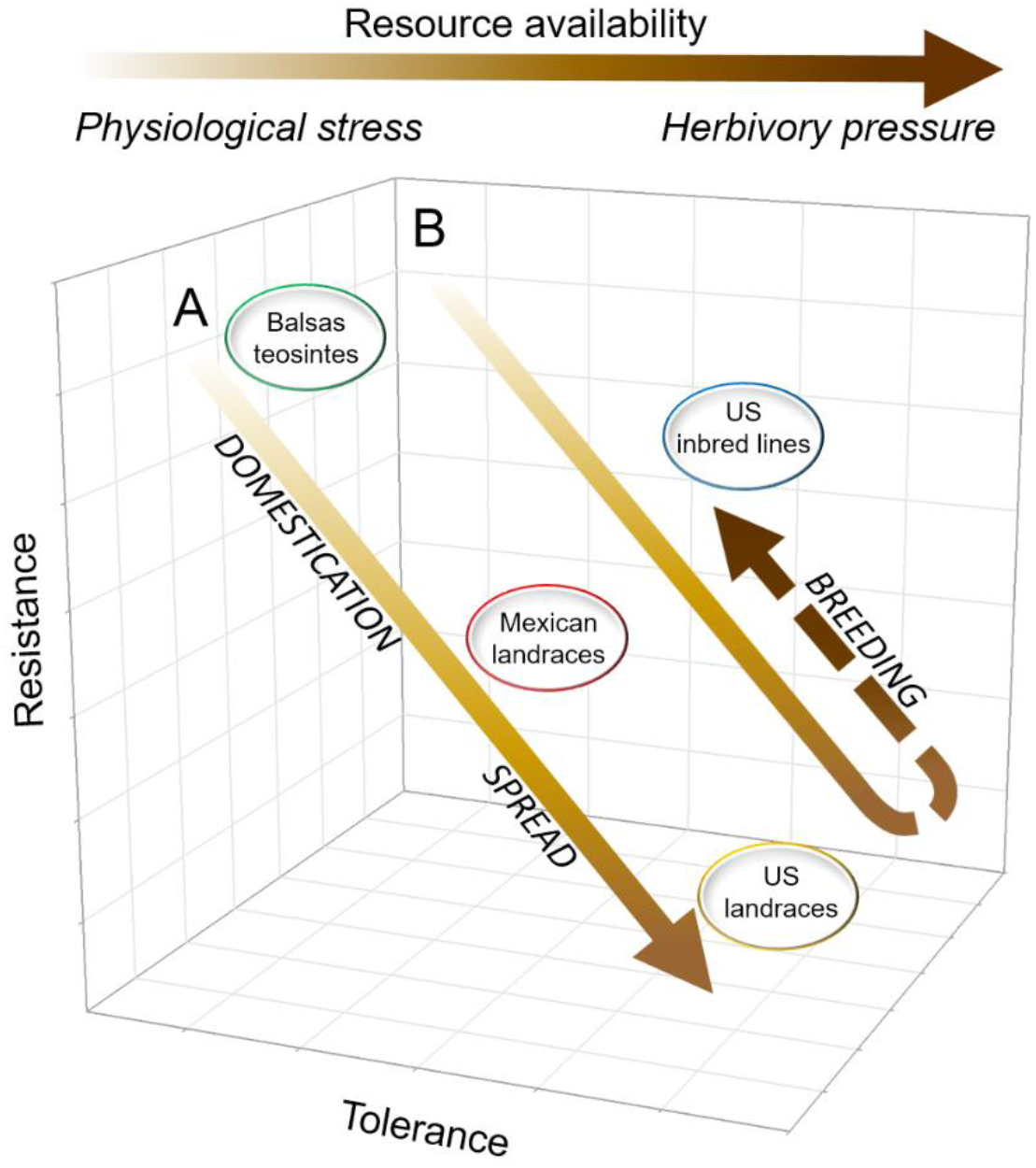
Hypothesized relationship between plant tolerance and resistance in maize, as mediated by agricultural intensification, resource availability, and environmental stress. In this study’s context, *Agricultural intensification* refers to widespread cultivation of high-yielding maize cultivars developed through systematic breeding, under the umbrella of chemical inputs, particularly commercial insecticides and synthetic fertilizers, and under increasing WCR pressure (see *Text* for additional details). The high-yielding cultivars are hybrids generated from inbred lines, which require chemical fertilizers (and adequate moisture) and pesticides to reach maximum productivity. The prior, *pre-intensification* period is characterized by widespread cultivation of landrace maize, natural and farmer (artificial) selection of landraces for broad resistance to environmental stresses, absence of fertilizers and pesticides, and minimal WCR pressure. **(A)** Prior to agricultural intensification, resistance to WCR gradually decreases while tolerance increases with maize domestication and spread, as resource availability increases, and as physiological stress gradually loses relevance to defense strategy evolution. **(B)** The trend of WCR resistance loss with WCR tolerance gain is reversed with breeding under agricultural intensification, where resource availability is high, physiological stress is minimized with the advent of chemical fertilizers, and WCR pressure becomes relevant to defense strategy evolution, though its relevance is mediated by insecticide use. In arrows in both **(A)** and **(B)**, and in horizontal arrow at top of figure, the lighter to darker gradient in coloration indicates an increasing gradient of resource availability; within this gradient, physiological stress and herbivory pressure are most relevant to defense strategy evolution at the low- and high-resource availability extremes, respectively.

## CONCLUSION

Undoubtedly, domestication was a consequential process that significantly affected maize growth, reproduction, and herbivore defense. Similarly, its spread from present-day Mexico exposed maize to new environments, and confronted it to novel suites of herbivores, which adopted the novel crop as a host because of its abundance and advantages, e.g., weakened defenses, superior nutritional value, refuge from natural enemies. With domestication and spread, the distribution and abundance of maize increased beyond those of Balsas teosinte, its wild ancestor, and with those increases maize broadened its genetic diversity as it was challenged by novel abiotic and biotic stresses (Hufford et al., 2012a; Hufford et al., 2012b; Bellon et al., 2018). Breeding in the last 100 years narrowed maize’s genetic diversity to increase its productivity in the context of resource-rich environments (including fertilizers, irrigation, and pesticides) in which tolerance- and resistance-based defenses were favored or neglected through systematic breeding, mainly for yield. In parallel, and partly as a consequence of the increasingly resource-rich environment in which maize was cultivated, WCR became a significant pest of maize in the US. As maize agriculture intensified beginning in the mid-1900s, it seems that tolerance as a basis for WCR management was neglected in deference to chemical control, though a small degree of WCR resistance was gained through breeding. Thus, it seems that US maize inbred lines, the parents of commercial hybrid varieties, are neither tolerant nor resistant to WCR, so are reliant on external means of defense against this pest, such as insecticides. Overall, our results suggested that the evolution of defense strategies in maize is predicted by ecological-evolutionary hypotheses seeking to explain defense strategy evolution in plants generally, within the contexts of plant resistance-productivity trade-offs, plant tolerance-resistance trade-offs, and varying resource availability vis-à-vis plant physiological stress and herbivory pressure.

## Author Contributions

AAFP and JSB contributed significantly to the conception, design, analysis, interpretation of data; drafted the article and revised it for important intellectual content. AAFP performed all experiments and collected the data. AAFP and JSB conducted statistical analyses and interpretation, and approved the final version to be published.

## Acknowledgments

We are grateful to M. J. Kolomiets and his laboratory (Texas A&M University, College Station), who kindly provided technical support and maize seeds. Dr Mark Millard (USDA NPGS, Ames, IA) provided Mo17, and Mexican and US maize landrace seed, and Chad Nielson (USDA ARS, North Central Agricultural Research Laboratory, Brookings, SD) provided Western corn rootworm eggs. M. J. Kolomiets, R. F. Medina, and K. Zhu-Salzman (all Texas A&M University, College Station) helped us improve an earlier version of the manuscript.

